# Capturing cell type-specific chromatin structural patterns by applying topic modeling to single-cell Hi-C data

**DOI:** 10.1101/534800

**Authors:** Hyeon-Jin Kim, Galip Gürkan Yardımcı, Giancarlo Bonora, Vijay Ramani, Jie Liu, Ruolan Qiu, Choli Lee, Jennifer Hesson, Carol B. Ware, Jay Shendure, Zhijun Duan, William Stafford Noble

## Abstract

Single-cell Hi-C (scHi-C) interrogates genome-wide chromatin interaction in individual cells, allowing us to gain insights into 3D genome organization. However, the extremely sparse nature of scHi-C data poses a significant barrier to analysis, limiting our ability to tease out hidden biological information. In this work, we approach this problem by applying topic modeling to scHi-C data. Topic modeling is well-suited for discovering latent topics in a collection of discrete data. For our analysis, we generate twelve different single-cell combinatorial indexed Hi-C (sciHi-C) libraries from five human cell lines (GM12878, H1Esc, HFF, IMR90, and HAP1), consisting over 25,000 cells. We demonstrate that topic modeling is able to successfully capture cell type differences from sciHi-C data in the form of “chromatin topics.” We further show enrichment of particular compartment structures associated with locus pairs in these topics.

## 1 Introduction

Single-cell chromosome conformation capture methods, such as single-cell Hi-C (scHi-C) [1–3], enable quantitative assessment of 3D conformation of chromosomes in individual cells. The resulting data can be used to detect cell-to-cell variations in genome-wide chromatin interactions and has the potential to interrogate chromosome structural heterogeneity in different cell types, thereby extracting important biological features that are otherwise hidden in bulk Hi-C data. However, the low coverage and consequent sparse nature of scHi-C data present a major computational barrier to the analysis. In each individual cell, scHi-C measures only a very small fraction of extant chromosomal contacts. Thus, to a much greater extent than in bulk Hi-C, the absence of an observed contact between two genomic loci does not provide strong evidence for the absence of a contact between those loci.

Currently, a limited number of methods have been developed for the analysis of scHi-C data. We previously developed a similarity-based embedding method called HiCRep/MDS to project scHi-C data into low dimensional space and arrange cells according to their cell-cycle phases [4]. This approach leverages a similarity measure, stratum adjusted correlation coefficient, that was developed for comparing bulk Hi-C matrices [5], combined with multidimensional scaling (MDS) to preserve distances between individual scHi-C contact maps. However, this method is computationally costly. Furthermore, although HiCRep/MDS can very accurately recover cell cycle information from several Hi-C data sets, the method frequently fails to capture chromatin structural differences between cell types [6].

More recently, Zhou *et al.* applied linear convolutions and random walks to scHi-C matrices to cluster cells according to their cell types [6]. This method uses imputation to reduce the sparsity of scHi-C matrices and identify chromatin structures that are similar to topologically associated domains within single cells. Despite the benefits this method offers, it lacks the ability to capture a set of genomic interactions that distinguish between cell types and to allow us to biologically interpret what features drove the clustering.

To overcome these limitations, we turned our attention to topic modeling. Topic modeling is mainly used in natural language processing to tackle challenging problems in text mining such as discovering hidden structures (or “topics”) in large collections of sparse, discrete data. One widely used approach to topic modeling is latent Dirichlet allocation (LDA), an unsupervised generative probabilistic model for a collection of documents [7]. In this setting, each document is represented as a vector of word frequencies. The LDA models this vector as a mixture of topics, where each topic is represented as a probabilistic distribution over words. LDA then assumes that a document is generated by the following steps: (1) choosing a topic mixture for the document according to a Dirichlet distribution over some specified number of topics, and (2) generating words in the document by iteratively sampling a topic from the topic mixture and selecting a word from the topic’s multinomial distribution [8]. With this generative model, the LDA identifies a set of topics that have most likely generated a given collection of documents. In the end, LDA generates two matrices, one that describes the relationship between topics and document and the other describing the relationship between topics and words [9]. In the first matrix, topics with the highest contributions are those that best characterize a document. In the second matrix, words with the highest contributions in a topic provide a sense of what the topic might represent.

Recently, González-Blas *et al.* applied LDA topic modeling to single-cell epigenomic data to simultaneously discover cell types and latent topics that pertain to topic-specific regulatory programs [10]. This approach applied topic modeling to single-cell ATAC-seq data to predict cell type-specific gene regulatory networks in the human brain.

Motivated by this work, we hypothesized that LDA could be employed to extract biologically meaningful structure from single-cell Hi-C data. In our setting, each cell corresponds to a document, and each locus pair corresponds to a word. We aimed to uncover features of chromatin structures—chromatin topics—that are characteristic of particular cell types. In our work, we generate single-cell Hi-C data from five human cell lines (GM12878, H1Esc, HFF, IMR90, and HAP1) using single-cell combinatorial indexed Hi-C (sciHi-C) [3] and show that applying topic modeling to sciHi-C data allows us to successfully cluster cells by cell types and discover cell type-specific topics of locus pairs. We demonstrate how the LDA identifies locus pairs that characterize each topic, and we show that the topics carry information about cell type-specific locus pair compartment structure.

## 2 Materials and methods

### 2.1 Data sets

Twelve human sciHi-C libraries were generated from five human cell lines (Table 1), which are available via the 4D Nucleome data portal (http://4dnucleome.org) [11].

**Table 1:**
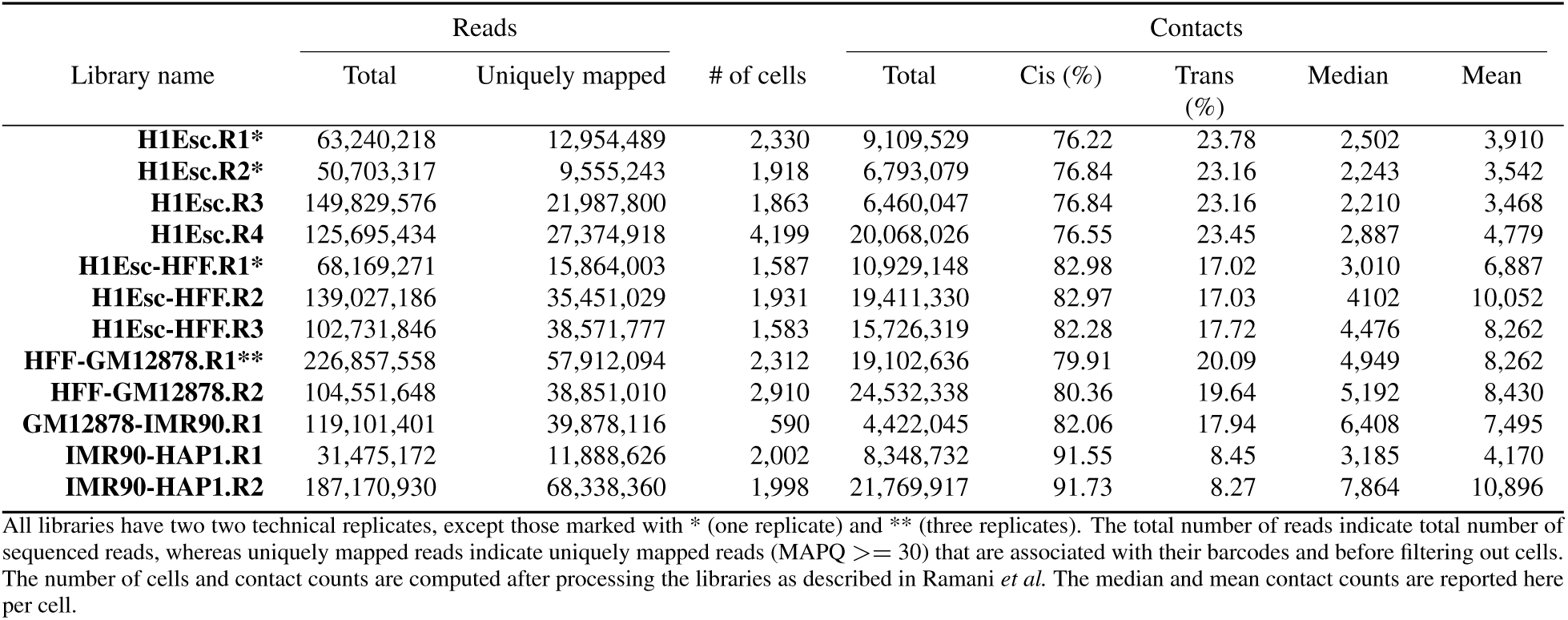
Summary statistics of all twelve sciHi-C libraries.

#### Cell culture

Human embryonic stem cells (hESC) H1, HFF-hTERT clone #6 (HFFc6), GM12878, IMR90 and HAP1 cells were obtained through the 4D Nucleome Consortium and cultured following the respective protocols standardized by the 4D Nucleome Consortium (4DN SOPs, https://www.4dnucleome.org/cell-lines.html).

#### SciHi-C library construction

SciHi-C libraries were generated as previously described in Ramani *et al.* [3] with some modifications. Briefly, cells grown at the exponential phase were fixed by incubating with formaldehyde at room temperature (25° C) for 10 min. The final concentration of formaldehyde was 2% for HFFc6, GM12878, IMR90 and HAP1 cells, and 3.5% for H1ESCs. Crosslinking was quenched by incubating with 0.125 M glycine on ice for 15 min. Crosslinked cells were then permeabilized with their nuclei intact and subjected to DpnII digestion to fragment the chromatin. Nuclei were then distributed to 96 wells, wherein the first-round barcodes were introduced through ligation of barcoded biotinylated double-stranded bridge adaptors. Intact nuclei were then pooled and subjected to proximity ligation, followed by dilution and redistribution to a second 96-well plate (no more than 25 nuclei per well). Crosslinking was reversed by incubating the nuclei in 2x NEB buffer 2 at 65° C overnight, followed by digestion with restriction enzymes Alu I and Hae III at 37° C overnight. After dA-tailing, second round barcodes were introduced through ligation of barcoded Y-adaptors. A total of 0.8 volume of Ampure beads were then added to each well, and all reaction samples in the 96-well plate were once again pooled and purified, and biotinylated junctions were then purified with Dynabeads M-280 streptavidin beads. Illumina sequencing libraries were amplified by PCR (13–15 cycles were used for each 96-well sample) and purified by two rounds of 0.8 volume Ampure beads. Libraries were sequenced by 2 × 250 bp paired-end run on a HiSeq-2500.

#### Mapping

Reads were processed according to the protocol described in Ramani *et al.* [3], mapping to human genome assembly hg19 using Bowtie2 2.2.3 [12] with default settings. As in previous work, cells with fewer than 1,000 unique reads, cis/trans-chromosomal contact ratio lower than 1, or uniquely mapped read percentage lower than 95% were filtered out. After this mapping step, the data covers 25,223 cells with a mean coverage of 6,608 sciHi-C contacts per cell. The sciHi-C contacts for each cell were aggregated into a square contact map using bins of 500 kb.

### 2.2 Topic modeling

To apply topic modeling to sciHi-C data, we adapted the protocol that González-Blas *et al.* proposed for single-cell RNA-seq and ATAC-seq data. As in their cisTopic model, we treated cells as documents, but in our case the role of words is played by genomic locus pairs (LPs). For our analysis, we only considered intra-chromosomal locus pairs that are within 10 Mb of one another. With sciHi-C data binned at 500 kb resolution and only considering the contacts in autosomal chromosomes (chr 1–22), this range includes 111,340 unique locus pairs. In our data, these locus pairs receive 68.7% of the observed contacts.

### 2.3 Method assessment

To evaluate the performance of topic modeling, we computed an AUC score from the cell-topic matrix for every cell. First, we reduced the cell-topic matrix with UMAP [13] and then calculated pairwise Euclidean distances between the cells in the UMAP space. For each cell, we binarized the cell type labels in our datasets (0 if correct cell type, 1 otherwise) and used the pairwise Euclidean distances as target scores and the resulting binarized cell type labels as true labels to compute an AUC score. We iterated this procedure for every cell in our datasets and computed an average AUC score for each cell type. For comparison, we computed average cell type-specific AUC scores from the UMAP reduced raw data.

### 2.4 Testing for topic specificity

The two-sample Wilcoxon test determines whether two empirical distribution functions significantly differ from one another [14]. We conducted one-sided, two-sample Wilcoxon tests on the cell-topic weight distributions from two different cell types to determine cell type specificity of topics. Since our datasets are comprised of five cell types, we conducted four separate pairwise Wilcoxon test for each cell type for each topic, thereby ensuring that the topic is specific to only the cell type we are comparing with. For example, if we were to test whether topic 1 is specific to GM12878, we would perform the one-sided Wilcoxon test on the topic 1 weight distributions of H1Esc and GM12878, HFF and GM12878, IMR90 and GM12878, and HAP1 and GM12878. If the Benjamini-Hochberg (BH) adjusted p-values for all four tests were below the significance level at 0.01, then the topic was assigned to that cell type.

## 3 Results

### 3.1 Applying topic modeling to sciHi-C datasets

We used topic modeling to decompose our sciHi-C data into a collection of topics (Figure 1). To do so, we represented the data in a binary matrix with 25,223 row (cells) and 111,340 columns (locus pairs) and used the matrix as input for LDA. To select the number of topics, we ran LDA over a range of numbers of topics, selecting the value 95 that yielded the maximum log-likelihood on held-out data (Supplementary Figure S1). The output of this analysis consists of two matrices: a 25,223 × 95 cell-topic matrix and a 111,340 × 95 LP-topic matrix (Supplementary Figure S3).

**Figure 1:**
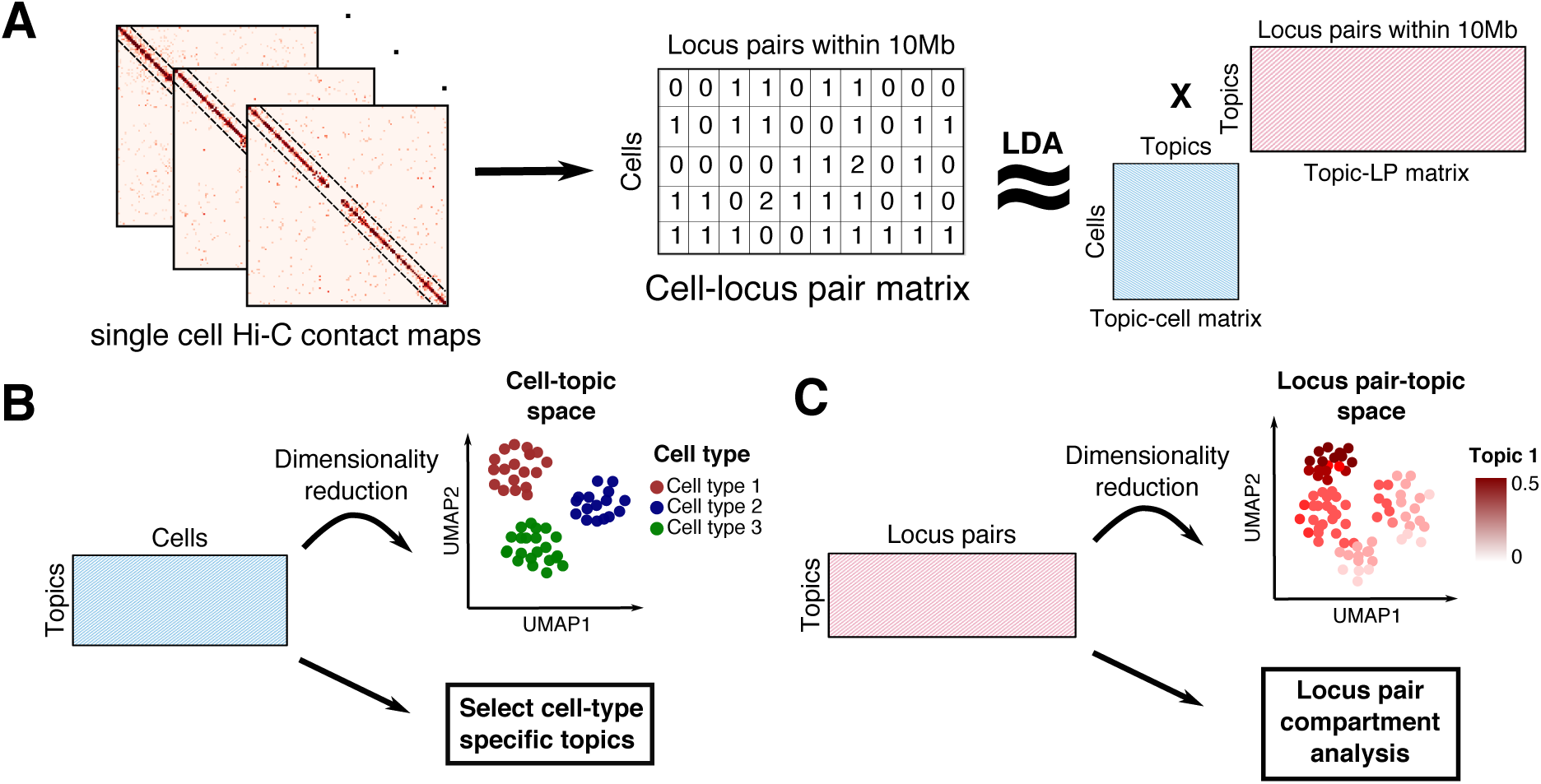
Topic modeling of sciHi-C data. (A) SciHi-C data was binned at 500kb resolution and processed into a cell-by-locus pair (LP) matrix. The resulting cell-LP matrix was provided as input to LDA to produce cell-topic and LP-topic matrices. Only intra-chromosomal contacts spanning distances <10 Mb were considered for analysis. (B) The output cell-topic matrix is projected into 2D using UMAP and can also be used to select cell type-specific topics. (C) The output LP-topic matrix can be used for selecting a subset of locus pairs that are most representative of each topic and for performing locus pair compartment comparison.

To evaluate whether topic modeling successfully extracted biological information from the sparse sciHi-C data, we applied Unifold Manifold Approximation and Projection (UMAP) [13] to the resulting cell-topic matrix (Figure 2A). Most notably, the data exhibits strong clustering of cells by their cell types, suggesting that cells of the same cell type have similar topic contribution profiles. An exception is HFF and IMR90 cells, which are embedded close together, likely reflecting the fact that they are both fibroblasts (foreskin and fetal lung, respectively). An additional cluster was observed in the embedding, composed of cells from all five cell types. The identity of this cluster is unknown, although the cells in the cluster exhibit high numbers of distinct contacts.

**Figure 2:**
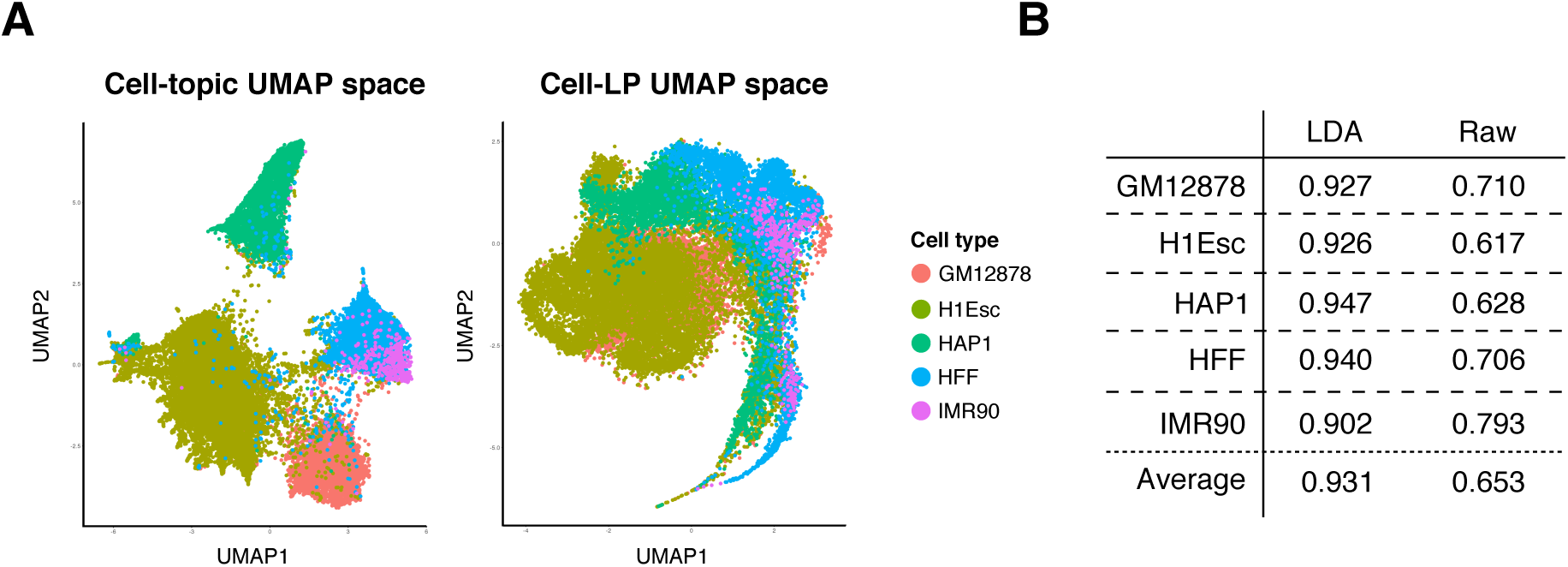
Comparison of topic reduced data and raw data. (A) The cell-topic matrix and raw data matrix were reduced to 2D with UMAP. (B) The UMAP projection coordinates and true cell type labels were used to compute an average AUC score for each cell type. The cell type-specific AUC scores computed with UMAP reduced cell-topic matrix are higher than that of UMAP reduced raw data across all cell types.

We also investigated the effect of potential confounders on the learned embedding. Visualization of the data with color according to sequencing library shows little evidence of batch effects (Supplementary Figure S2B). Similarly, repeating the LDA using data that was downsampled led to a similar separation of cell types in the 2D UMAP space (Supplementary Figure S5A). Specifically, downsampling was done by computing the median contact count (2,432 contacts) across the 111,340 proximal locus pairs, discarding cells with fewer than the median number of contact, and randomly downsampling the remaining 12,613 cells to median number of contacts. Within each cell type, we observe some clustering by coverage (Supplementary Figure S2C), suggesting that the effect of coverage on the learned model is small relative to the differences between cell types.

### 3.2 Evaluating the performance of topic modeling on cell type clustering

We evaluated the performance of topic modeling on clustering based on cell types. To do so, we computed cell type-specific AUC scores by using the UMAP projected cell-topic matrix and their cell type identities. For each cell in the UMAP space, we first calculated pairwise Euclidean distances between all the other cells. We then binarized the cell type labels in the dataset relative to other cells (0 if same cell type, 1 otherwise) and used the binarized labels and Euclidean distances to compute an AUC score for that cell. We iterated this process for every cell and took the average scores across each cell type for evaluation.

Applying topic modeling to our datasets significantly improved the clustering of cells by their cell types, with an average AUC score of 0.931 vs 0.653 computed from UMAP projected raw data (Figure 2B). The average cell type-specific AUC scores computed from the cell-topic matrix were higher in all cases, suggesting that topics contain cell type-specific information. We attempted to use other embedding methods, such as HiCRep/MDS and scHiCluster on our datasets for comparison, but we were not able to do so with reasonable computational resources, mainly because these methods require all intrachromosomal pairwise contact matrices from 25,223 cells. On the other hand, we were able to perform topic modeling on our cell-LP matrix in a day by using several CPUs. Altogether, the superior performance of topic modeling on cell type clustering relative to the raw data along with relatively cheap computational costs highlight the advantages of applying topic modeling to single cell Hi-C data.

### 3.3 Identifying and characterizing cell type-specific topics

Visualization of the cell-topic matrix (Figure 2A) suggested that many of the topics are highly associated with particular subpopulations of cells. Accordingly, to better understand this topical structure, we used a Wilcoxon test to identify topics that are specifically enriched for each of the five cell types (Section 2.4). This test assigned a total of 23, 22, 13, 11, and 4 topics specific to GM12878, H1Esc, HAP1, HFF, and IMR90, respectively.

#### 3.3.1 Examining topic associated locus pair compartment structure within cell type

To understand what each cell type-specific topic captures in the sciHi-C data, we evaluated the Hi-C compartments of the locus pairs that characterize each topic. Previous analysis of bulk Hi-C data suggests that chromatin can be usefully segregated into two compartments, A and B, where the A compartment is associated with euchromatin and the B compartment with heterochromatin [15]. Because these compartment structures vary by cell type [15, 16], we hypothesized that cell type-specific topics might be enriched for locus-pairs the exhibit cell type-specific compartment assignments.

Our compartment analysis consists of three steps: (1) assigning compartment labels to LPs in a cell type-specific fashion, (2) assigning topic labels to LPs based on the LP-topic matrix, and (3) evaluating whether a particular locus pair compartment structure is enriched in each cell type-specific topic.

For the first step, we used the aggregated sciHi-C data to assign compartments to GM12878, H1Esc, and HFF at a resolution of 500kb. IMR90 and HAP1 were excluded from this analysis due to low coverage of the cells. Using these compartment calls, each locus-pair was assigned one of three labels in each cell type, based on the cell type-specific compartment assignments at the midpoint of each locus: (1) both loci are active (AA), (2) both loci are inactive (BB), or (3) one locus is active and the other inactive (AB) (Figure 3A).

**Figure 3:**
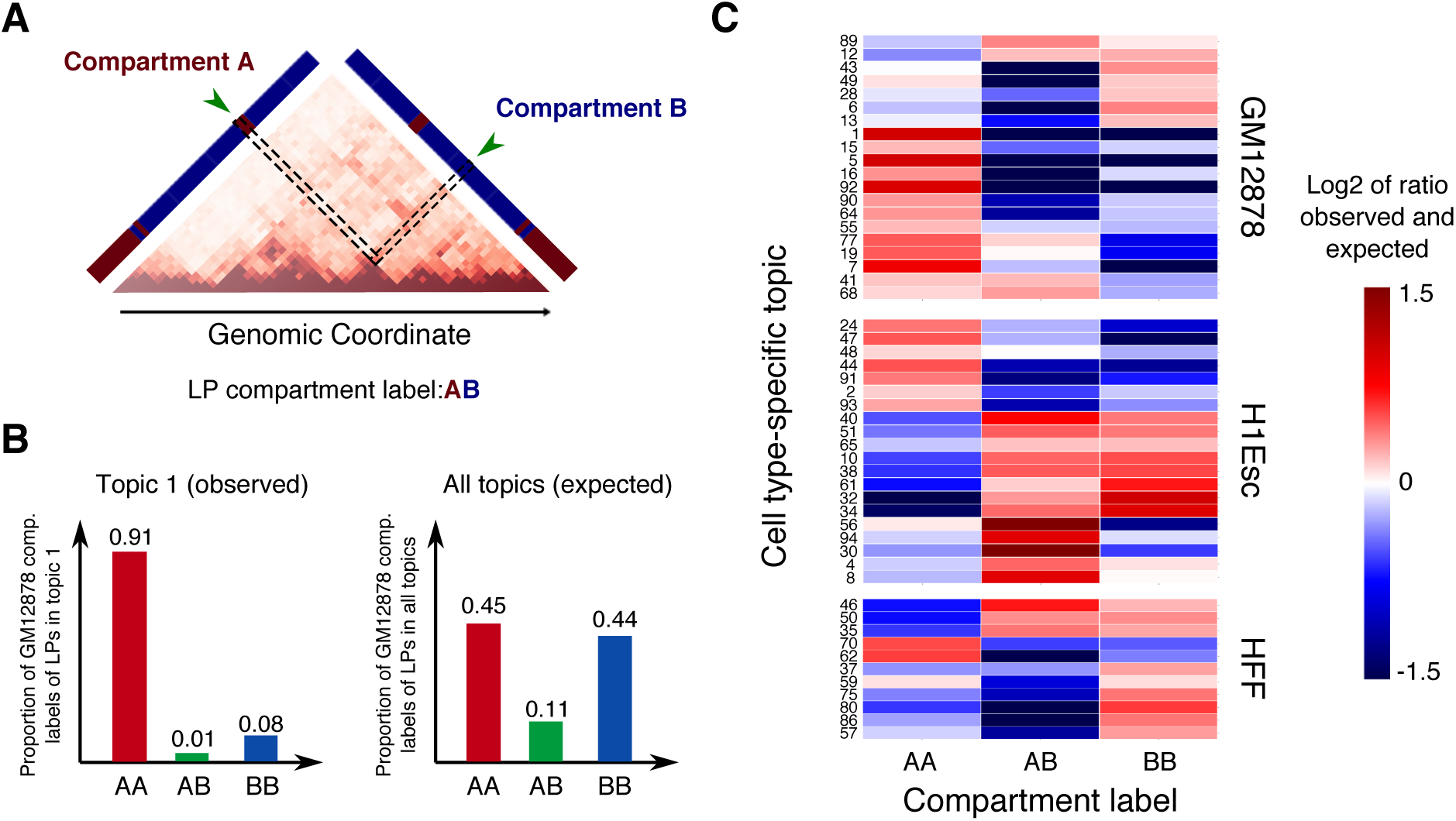
Analysis of compartment calls for cell type-specific topic associated locus pairs within cell type. (A) Each locus pair is assigned a pairwise compartment call. As an example, the locus pair enclosed in the a black box is assigned “AB” in GM12878. (B) The proportions of GM12878 compartment labels were computed for LPs in topic 1, which is a GM12878 specific topic (left), and for LPs in all topics to estimate expected proportions (right). (C) A heatmap showing the log_2_ ratio of observed over expected proportion of each LP compartment structure in topics specific to GM12878, H1Esc, and HFF. The figure only includes topics whose enrichments are deemed significant at *α* = 0.01 according to a chi-square test after BH correction.

For the second step, a visualization of the LP-topic matrix suggested that a small number of locus pairs contribute strongly to each topic (Supplementary Figure S3B). Accordingly, we identified topic-specific LPs as the top 1% of locus pairs based on the topic weights.

Finally, we computed the observed proportion of each compartment label among LPs associated with each cell type-specific topic. To ascertain which particular compartment structure is enriched in each topic, we estimated the expected proportion of each compartment using the compartment labels of all topic-associated LPs. We then compared the observed proportions with the expected to measure enrichment of LP compartment structures (Figure 3B).

Our compartment analysis showed that each cell type-specific topic contains LPs that are enriched for a distinct compartment structure. We assessed the significance of differences in the expected and observed proportions using a chi-square test for independence at the significance level of 0.01 with BH FDR correction. From the list of topics that were found to be specific to GM12878, H1Esc, and HFF, the chi-square test identified a total of 20, 20, and 11 topics, respectively, that exhibit significantly different LP compartment composition from the expected. When we examined the locus pairs in each cell type-specific topic, we observed varying patterns of enrichment in compartment structure. Specifically, more than half of the GM12878-specific topics were enriched for the AA compartment and around third were enriched for the BB compartment, whereas in H1Esc more than half of the topics were highly enriched for both the AB and BB compartments (Figure 3C). Conversely, HFF-specific topics showed enrichment for the BB compartment. Overall, these observations support our hypothesis that topics can capture cell type-specific locus pair compartment information.

#### 3.3.2 Investigating differences in topic-associated locus pair compartment assignments between cell types

To further explore this idea, we next compared locus pair compartment structures across cell types. In this analysis, a locus pair is assigned three compartment labels, one label for each cell type (Figure 4A–B). To pursue the hypothesis that cell type differences may be captured in LPs of cell type-specific topics, we assessed cell type compartment differences for LPs that are associated with each cell type-specific topic.

**Figure 4:**
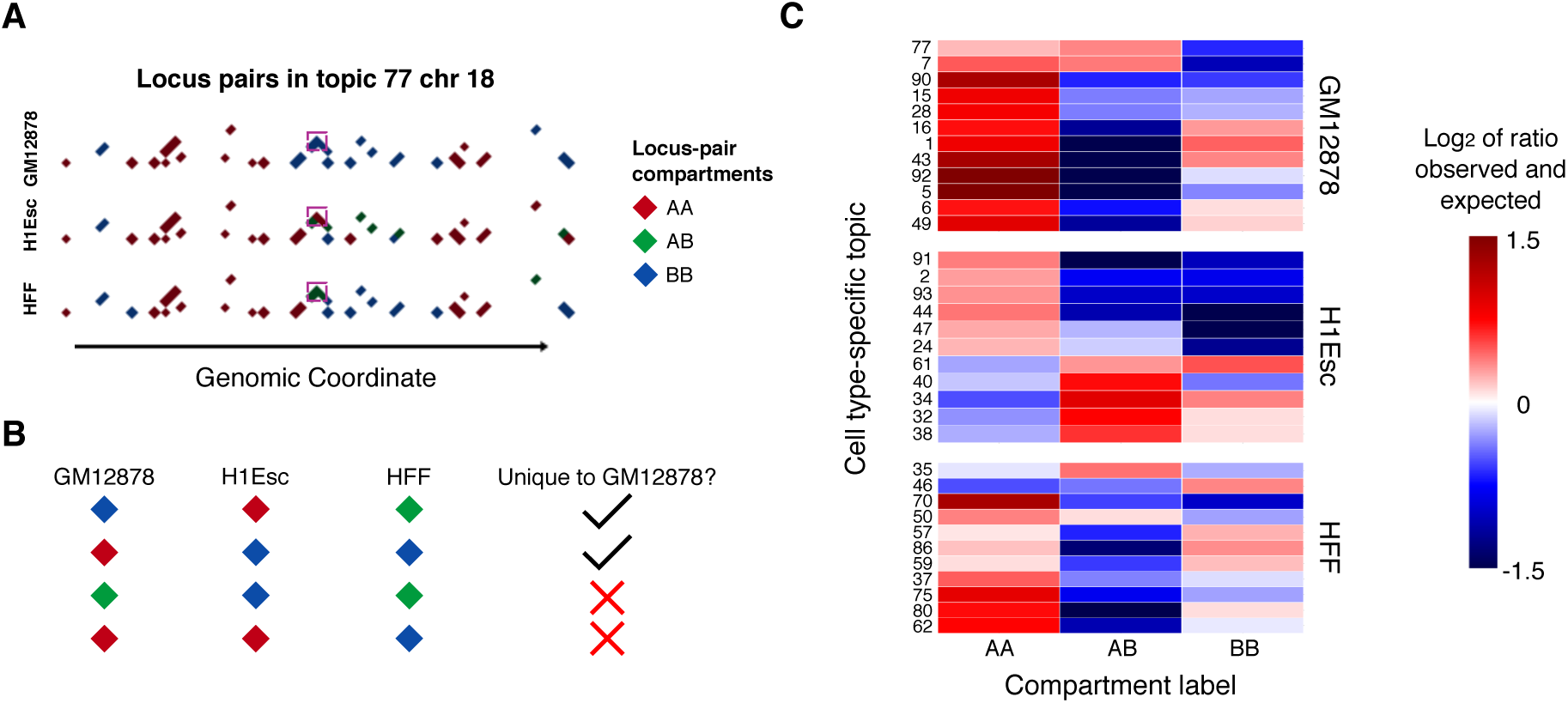
Quantifying differences in compartment assignment of topic-associated locus pairs between cell types. (A) A locus pair is tri-labeled, one label for each cell type. Here, the chr 18 locus pairs in topic 77 (GM12878 specific) were visualized for GM12878, H1Esc, and HFF, colored by compartment labels. The purple box highlights a locus pair that exhibits different compartment configurations between cell types. (B) Examples of LP compartment comparison between cell types. The first two rows show locus pairs that are unique to GM12878, whereas the last two rows show locus pairs that are not. (C) A heatmap showing the log_2_ ratio of observed over expected proportion of each cell type-specific compartment structure in topics specific to GM12878, H1Esc, and HFF. The figure only includes topics whose enrichments are deemed significant at *α* = 0.01 according to a chi-square test with BH FDR control.

First, for each set of cell type-specific topic LPs, we assigned three sets of compartment labels, corresponding to GM12878, H1Esc, and HFF. This yields a tri-label for each LP. We then compared the compartment labels between cell types and computed the total number of each compartment type (AA, AB, and BB) that is observed to be unique in the assigned cell type. For example, if a locus pair that was associated with GM12878-specific topic was assigned AA in GM12878 and BB in both H1Esc and HFF, we counted this locus pair as unique to GM12878 (Figure 4B). Using this approach, we computed the proportion of each compartment label that is unique to the assigned cell type for LPs in each topic.

In a similar fashion to Section 3.3.1, we then estimated the expected proportion of cell type-specific LP compartment labels using all topic-associated LPs, to measure the enrichment of a particular compartment structure in each cell type. We performed a chi-square test (*α* = 0.01, with BH correction) to determine the significance of the differences in observed and expected proportions. This analysis identified similar but fewer topics compared to the previous analysis: 12, 11, and 11 topics for GM12878, H1Esc, and HFF, respectively.

Overall, we observed an over-representation of particular compartment structures when the compartment labels of topic-associated LPs were compared between cell types. The similar pattern of enrichment was observed for LPs in H1Esc-specific topics as previously, in which LPs were more AB in H1Esc compared to other cell types (Figure 4C). Interestingly, the compartment labels of locus pairs that are unique to GM12878 and HFF were mainly enriched for the AA compartment, in contrast to the pattern observed in the previous compartment analysis. Indeed, when we took the set of LPs that are AA in each of the topics specific to GM12878 and HFF and computed the proportion of compartment labels in the other cell types, we observed a higher proportion of LPs that are in the AB or BB compartments than expected (Supplementary Figure S4). We obtained similar results when we performed the same analysis with the downsampled data (Supplementary Figure S5B–C). These observations suggest that active chromatin interactions (both LPs in the A compartment) are largely captured by cell type-specific topics, just as cisTopic was able to distinguish cell types from the single-cell ATAC-seq data by detecting cell type-specific accessible regions of chromatin [10].

## 4 Discussion

We have demonstrated that topic modeling of sciHi-C data identifies topics that successfully recapitulate cell type structure in the data, and that the topics themselves exhibit statistically significant patterns of enrichment relative to chromatin compartments. The decomposition of the sciHi-C data into the cell-topic and LP-topic matrices enables visualization of structure that is not otherwise apparent in the data. This decomposition also facilitates investigation of the “meaning” of topics relative to features such as chromatin compartment structure.

Topic modeling offers several important benefits in the context of sciHi-C data. The method was specifically developed to operate on sparse data such as sciHi-C data. The topics themselves potentially provides clues about which locus pairs are important in driving differences among cell types. In contrast, the existing single-cell Hi-C analysis methods HiCRep/MDS and scHiCluster mainly provide an embedding of the data, without offering the topic structure, and are much more computationally expensive than the topic modeling approach.

In future, we plan to investigate alternative ways to interpret the topics produced by our model. In our analysis, we investigated whether topics are associated with cell type-specific locus pair compartment structures. A more complete understanding of this structure may require also investigating finer-scale chromatin properties, such as topologically associating domain structure, or complementary genomic or epigenomic measurements of transcription, factor binding, or histone modifications. Such analysis is challenging because of the necessarily coarse scale (500 kb bin size in this work) of the topic modeling. Ideally, statistical procedures that jointly take into account sciHi-C data alongside bulk Hi-C and complementary linear genomic measurements are required to fully understand the relationships between 3D chromatin structure and fundamental cellular processes such as transcription and replication.

## Acknowledgments

This work was supported by National Institutes of Health award U54 DK107979. The funders had no role in study design, data collection and analysis, decision to publish, or preparation of the manuscript.

## 5 Supplemental Information

**Figure S1:**
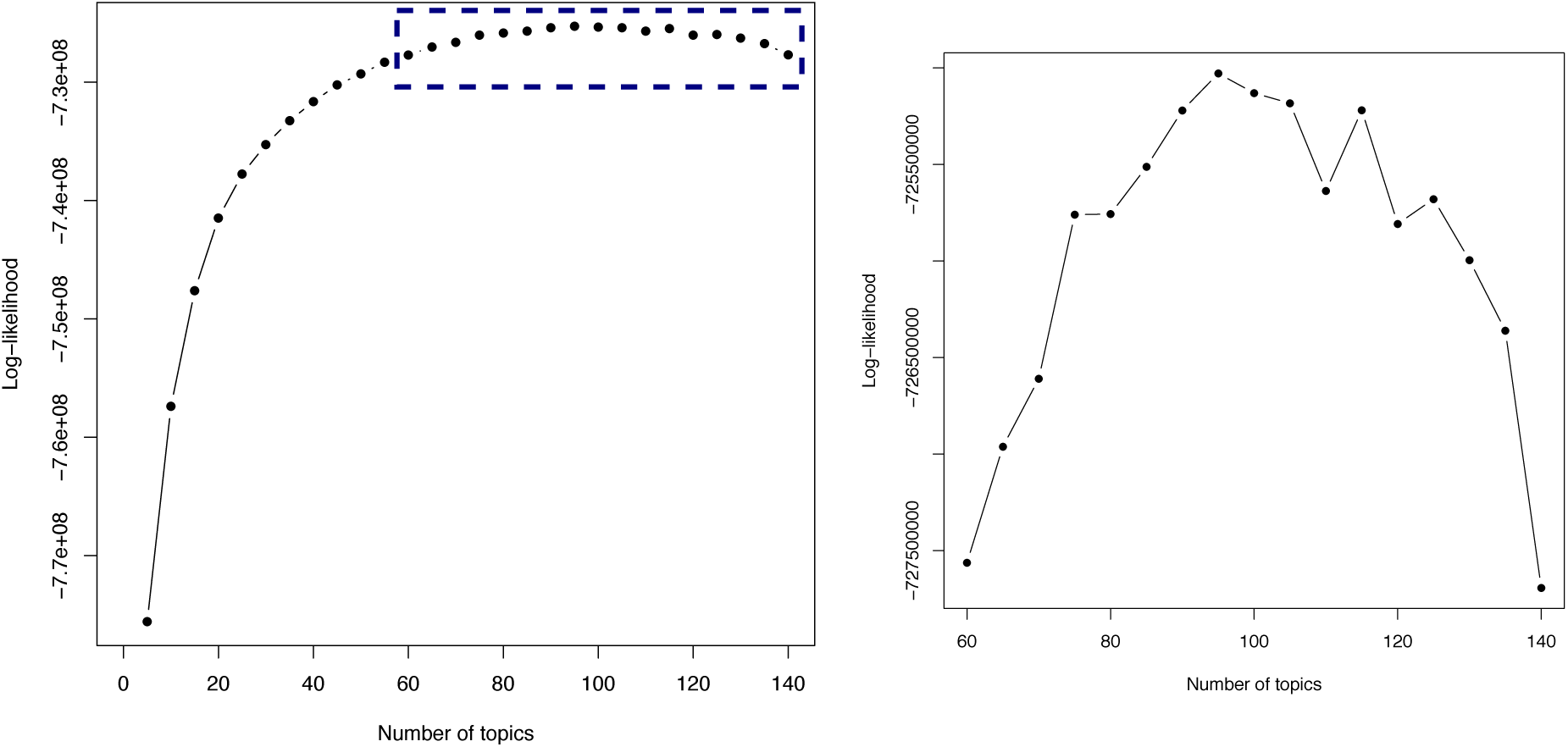
Selecting the number of topics. (Left) The figure plots the log-likelihood value at the last iteration of training as a function of the number of topics (5–140). (Right) The boxed portion of the left plot is zoomed in to highlight the highest log-likelihood at 95 topics.

**Figure S2:**
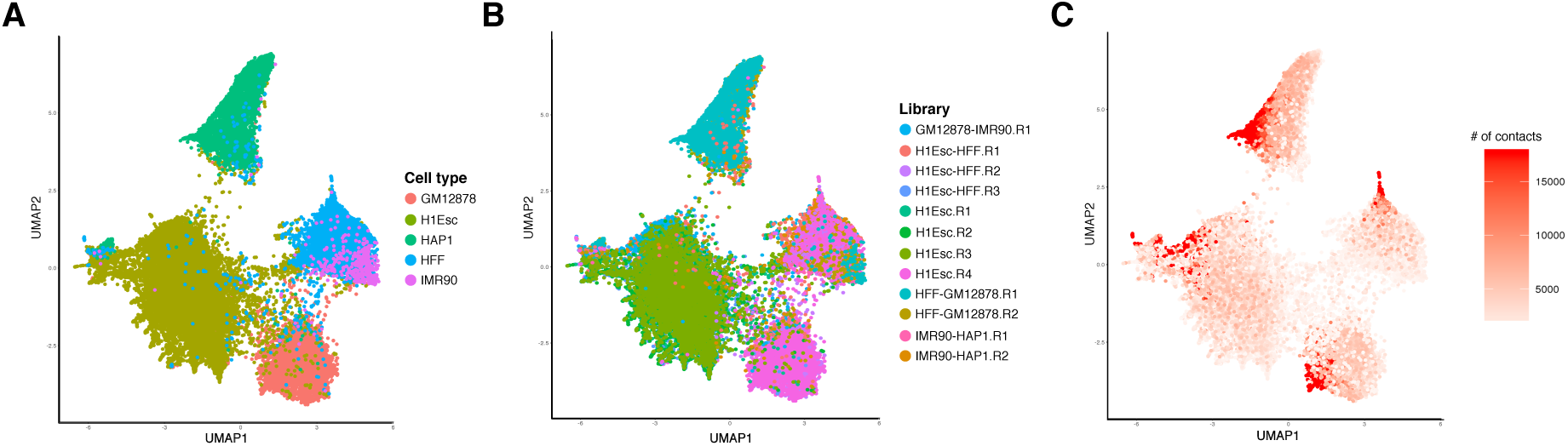
Topic modeling results shows little batch effects. (A) The 2D UMAP projection of cell-topic matrix from Figure 2A. The same plot colored by library (B) and (C) by coverage.

**Figure S3:**
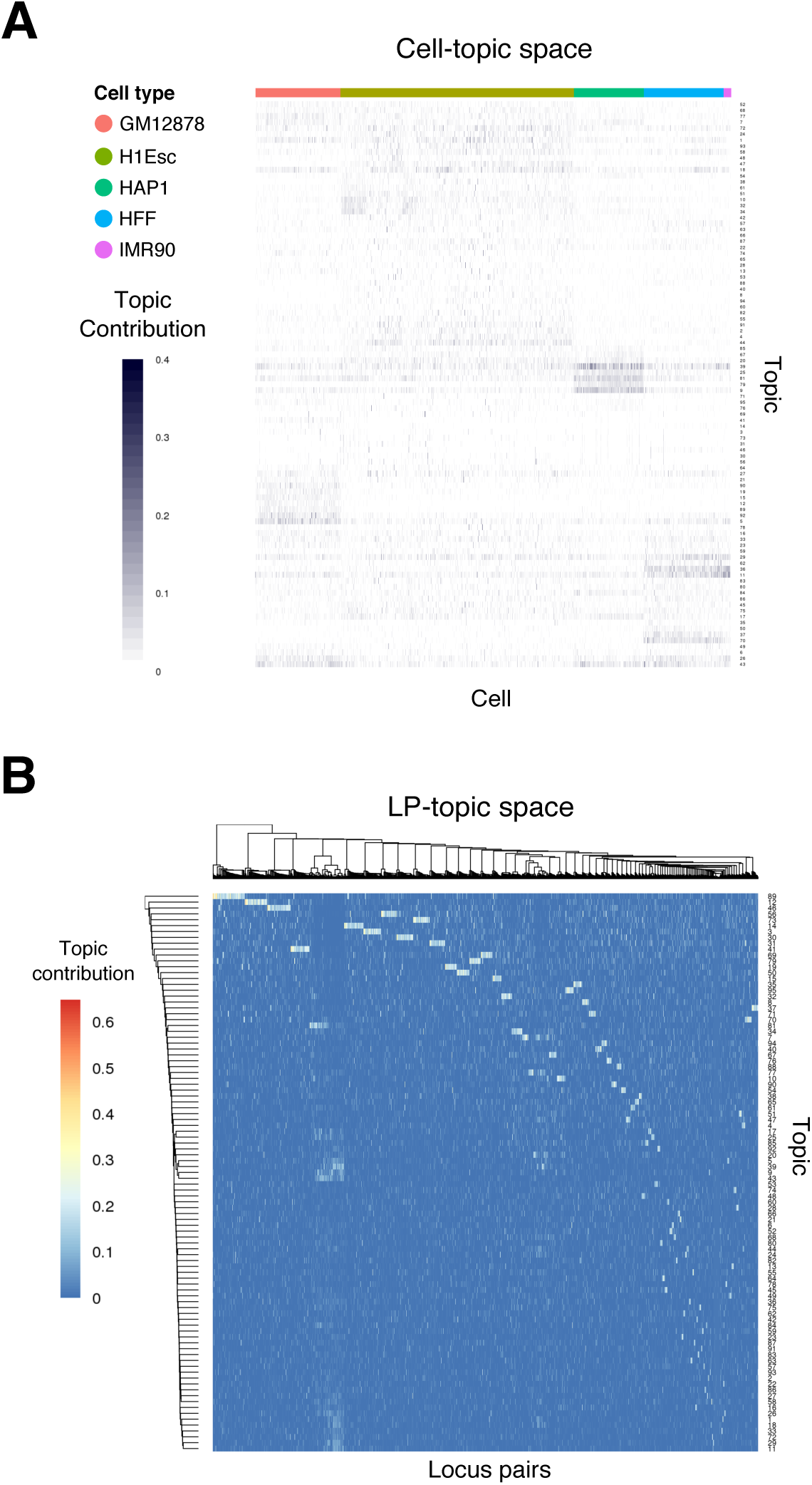
Visualization of cell-topic and LP-topic matrices. The resulting (A) cell-topic and (B) LP-topic matrices from topic modeling were visualized as heatmaps, with color bars indicating the value of topic weights. Topics are highly associated with particular subpopulations of cells and particular subset of locus pairs are associated with a topic.

**Figure S4:**
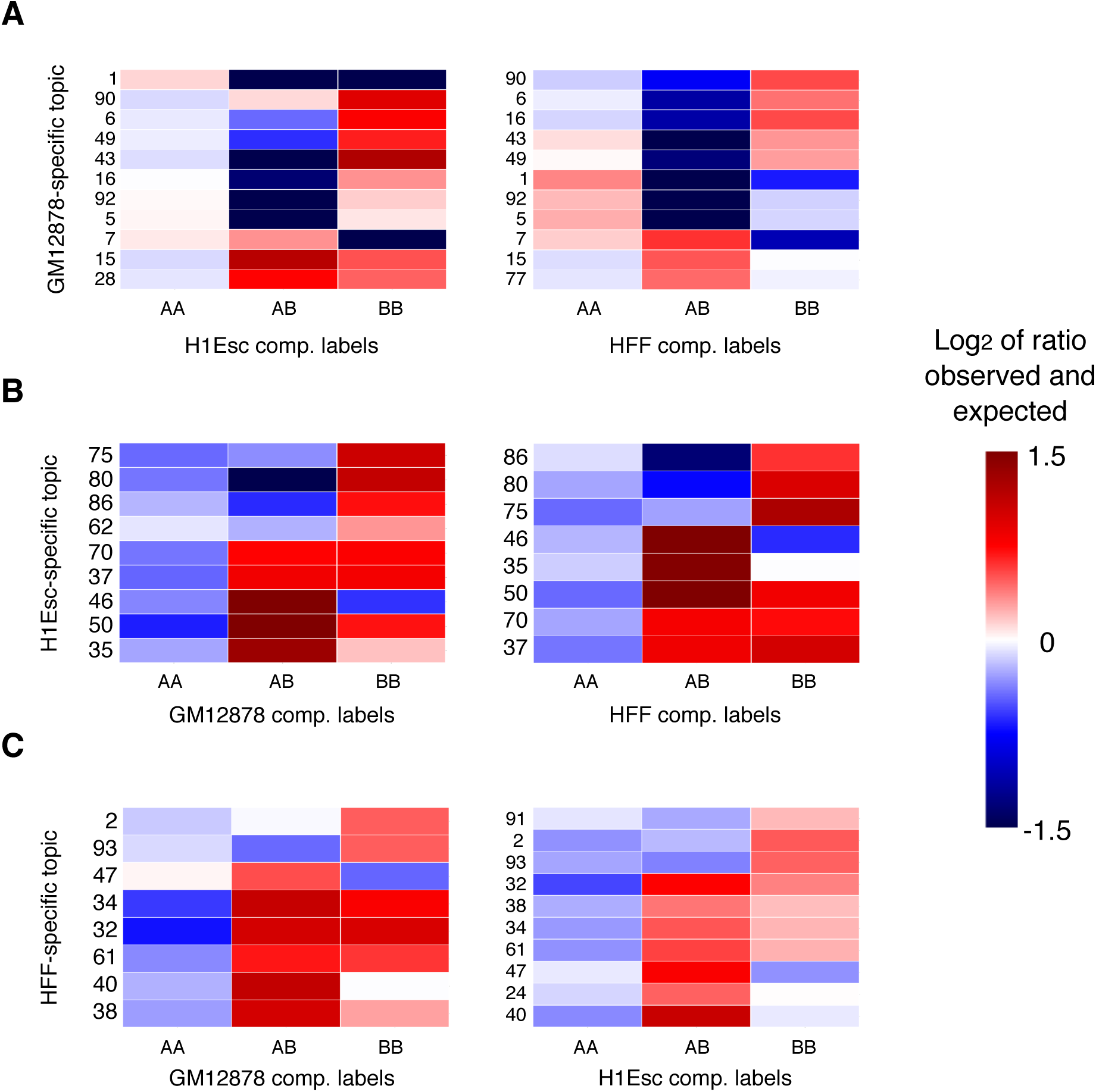
Analysis of compartment labels of AA LPs in the cell type-specific topics. For each GM12878-, H1Esc-, and HFF-specific topic, the locus pairs that are AA in the assigned cell type were assigned compartment labels in the other two cell types and their proportions were computed. Similar to the main compartment analysis, the observed proportions were compared with the expected proportions obtained from LPs in all topics. The comparison was done pairwise and enrichment was calculated for (A) H1Esc and HFF compartment labels of AA LPs in GM12878-specific topics, (B) GM12878 and HFF compartment labels of AA LPs in H1Esc-specific topics, and (C) GM12878 and H1Esc compartment labels of AA LPs in HFF-specific topics. The figure only includes topics whose enrichments are deemed significant at *α* = 0.01 according to a chi-square test with BH FDR control.

**Figure S5:**
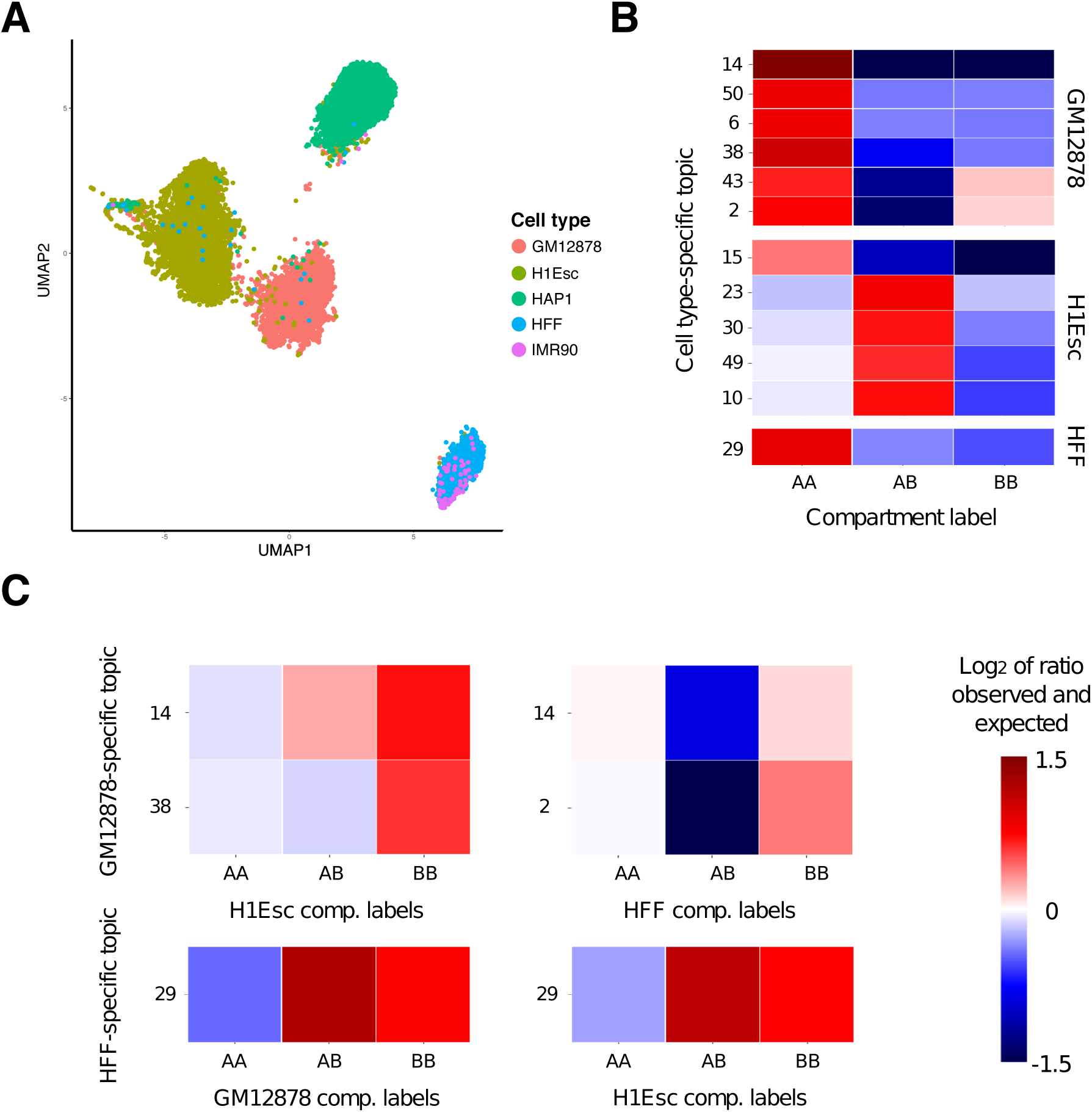
Analysis of topic modeling results using downsampled data. We computed the median number of contact in our data, discarded cells with fewer than this median number of contacts, and randomly downsampled the remaining 12,613 cells to the median. We applied LDA to the downsampled data and visualized the (A) cell-topic matrix in UMAP space, carried out compartment analysis to evaluate (B) topic-wise LP compartment assignment, and (C) AA LP compartment assignment differences between cell types. The number of topics yielded the maximum log-likelihood was 50, and the number of topics that were specific to GM12878, H1Esc, HAP1, HFF, and IMR90 were 12, 15, 4, 4, and 4, respectively.

